# The origin of extracellular DNA in bacterial biofilm infections *in vivo*

**DOI:** 10.1101/689067

**Authors:** Maria Alhede, Morten Alhede, Klaus Qvortrup, Kasper Nørskov Kragh, Peter Østrup Jensen, Philip Shook Stewart, Thomas Bjarnsholt

## Abstract

Extracellular DNA (eDNA) plays an important role in both the aggregation of bacteria and in the interaction of the resulting biofilms with polymorphonuclear leukocytes (PMNs) during an inflammatory response. Here, transmission electron and confocal scanning laser microscopy were used to examine the interaction between biofilms of *Pseudomonas aeruginosa* and PMNs in a murine implant model and in lung tissue from chronically infected cystic fibrosis patients. PNA FISH, DNA staining, labeling of PMN DNA with a thymidine analogue, and immunohistochemistry were applied to localize bacteria, eDNA, PMN-derived eDNA, PMN-derived histone H3 (H3), neutrophil elastase (NE), and citrullinated H3 (citH3). Host-derived eDNA was observed surrounding bacterial aggregates but not within the biofilms. H3 localized to the lining of biofilms while NE was found throughout biofilms. CitH3, a marker for neutrophil extracellular traps (NETs) was detected only sporadically indicating that most host-derived eDNA *in vivo* was not a result of NETosis. Together these observations show that, in these *in vivo* biofilm infections with *P. aeruginosa*, the majority of eDNA is found external to the biofilm and derives from the host.

**Author summary:** The role of extracellular DNA (eDNA) has been described *in vitro* to play a major role in biofilm formation and antibiotic tolerance, but never for biofilm infections *in vivo*.

Two important characteristics of human chronic bacterial infections are aggregated bacteria and white blood cells (WBC). Bacteria use eDNA to stabilize biofilms and WBC use eDNA to trap bacteria. Given the importance of eDNA for both bacteria and WBC we show here for the first time that bacterial biofilms do not co-localize with either bacterial or WBC-derived eDNA during chronic infections. Our *in vivo* findings show eDNA is located outside biofilms as opposed to incorporated within the biofilm as commonly observed *in vitro*.

We believe that understanding the interplay between biofilms and WBC during chronic infection is essential if we are to elucidate the mechanisms underlying persistent infection and ascertain why WBC fail to eradicate bacteria; this will enable us to develop new treatment strategies.

## Introduction

Persistent bacterial infections present an increasing challenge to health care practitioners worldwide. Aggregated bacteria, often referred to as biofilms, are tolerant and resistant to both antibiotics and the host immune system [1, 2]. Most studies of bacteria are based on planktonic cultures consisting primarily of single cells grown *in vitro* where both aggregates and host factors are missing [3]. To understand the underlying mechanism(s) causing persistence of chronic bacterial infections, it is important to investigate the interplay between aggregated bacteria and the host immune system. A common denominator of both bacterial biofilms and cells of the innate host defense is extracellular DNA (eDNA). *In vitro* studies have shown eDNA, produced by the bacteria themselves, to be important primarily for initial biofilm formation as a major structural component of bacterial biofilms [4] and as a stabilizer that contributes to increased tolerance of biofilms to antibiotics *in vitro* [5-8]. The common feature of chronic bacterial infections is persistent inflammation dominated by polymorphonuclear leukocytes (PMNs), which border bacterial aggregates but are unable to eradicate them [9]. This type of inflammation is classified as acute inflammation. Successful resolution of acute inflammation involves apoptosis and engulfment of demised PMNs by macrophages [10]. However, during chronic bacterial infections, PMNs are thought to undergo necrosis, during which release of toxic compounds such as NE, oxidants, and eDNA actually increase inflammation [10]. Furthermore, components such as NE contribute to the destruction of tissue in chronic wounds and at sites of inflammation in the lungs of cystic fibrosis (CF) patients [11-15]. PMNs actively secrete neutrophil elastase (NE) and histones attached to DNA, via a process termed NETosis [16], both *in vitro* and *in vivo* in acute infections [17, 18]. This process is collectively referred to as neutrophil extracellular traps (NETs), which capture and kill bacteria. Since 2004, formation of NETs has been proposed as a function of neutrophils and other immune cells [19] activated *in vitro* and *in vivo* in response to various bacterial components [18, 20]. However, a recent review questioned whether NETs play a role in innate immunity and thereby trapping of bacteria [21]. Release of eDNA by NETosis has, to the best of our knowledge, never been demonstrated in association with inflammation caused by chronic bacterial infections, despite evidence of close contact between PMNs and bacteria.

At present, our knowledge about the interaction between *in vivo* biofilms and the host immune system during chronic infections is limited, and we do not know the role or localization of eDNA in relation to the biofilms *in vivo*. Unsuccessful eradication of infectious biofilms despite accumulation of PMNs has not been explained fully. Thus, to obtain more detailed information about the interplay between bacterial aggregates and PMNs during chronic infection, and to increase our understanding of the role of eDNA, we studied these phenomena for the first time, directly in a murine implant model of biofilm infection and directly in human samples of chronic infected CF patients. By using transmission electron microscopy (TEM) we were able to study the interaction between PMNs and biofilm of *Pseudomonas aeruginosa* formed on a silicone implant inserted in the peritoneal cavity of mice. Furthermore, labeling the DNA of PMNs during their S-phase using the Click-iT technology *in vivo* and confocal scanning laser microscopy (CSLM), showed PMN-derived DNA did not localize with biofilms. To establish the localization of specific components originating from PMNs we used immunohistochemistry and found that PMN-derived histone H3 (H3) and neutrophil elastase (NE) co-localized with biofilms, but citrullinated H3 (citH3) and PMN-derived DNA did not. The results obtained using the mouse model could be correlated directly to chronic bacterial infections in human CF lungs.

## Results

### TEM-based examination of the interaction between biofilms and PMNs in a murine implant model shows damaged PMNs

Previously, we used scanning electron microscopy (SEM) [22] to examine the interaction between immune cells and *P. aeruginosa* biofilms in a murine implant model and found a marked influx of PMNs toward the biofilms. We then used CSLM to identify PMNs according to their segmented lobular nucleus. Here, we used TEM to examine the in-depth additional detailed interaction between bacteria and PMNs, and to visualize the matrix of the biofilm. Implants were inspected at three different time points (6 h, 24 h, and 48 h post-insertion) to illustrate progression of infection and the immune response.

At 6 h post-insertion, we noted intact PMNs containing internalized bacteria within vacuoles, which indicates active phagocytosis (Fig 1A). PMNs with internalized bacteria were confined to areas in which bacteria had not yet aggregated. No intact PMNs were observed in areas of high bacterial density (Fig 1B). At 24 h post-insertion, bacterial aggregation resulted in the formation of large biofilms lining the inside of the implants. PMNs frequently adjoined the bacteria. These PMNs were enlarged and had damaged cell membranes (Fig 1C). Intact PMNs at this time point were rare. Matrix material surrounding the bacteria within the biofilm was evident (Fig 1 E-G). At 48 h post-insertion, the biofilm appeared denser and the PMNs were tightly interfaced with the biofilms (Fig 1D). Matrix material was clearly visible (Fig S1).

**Fig. 1.**
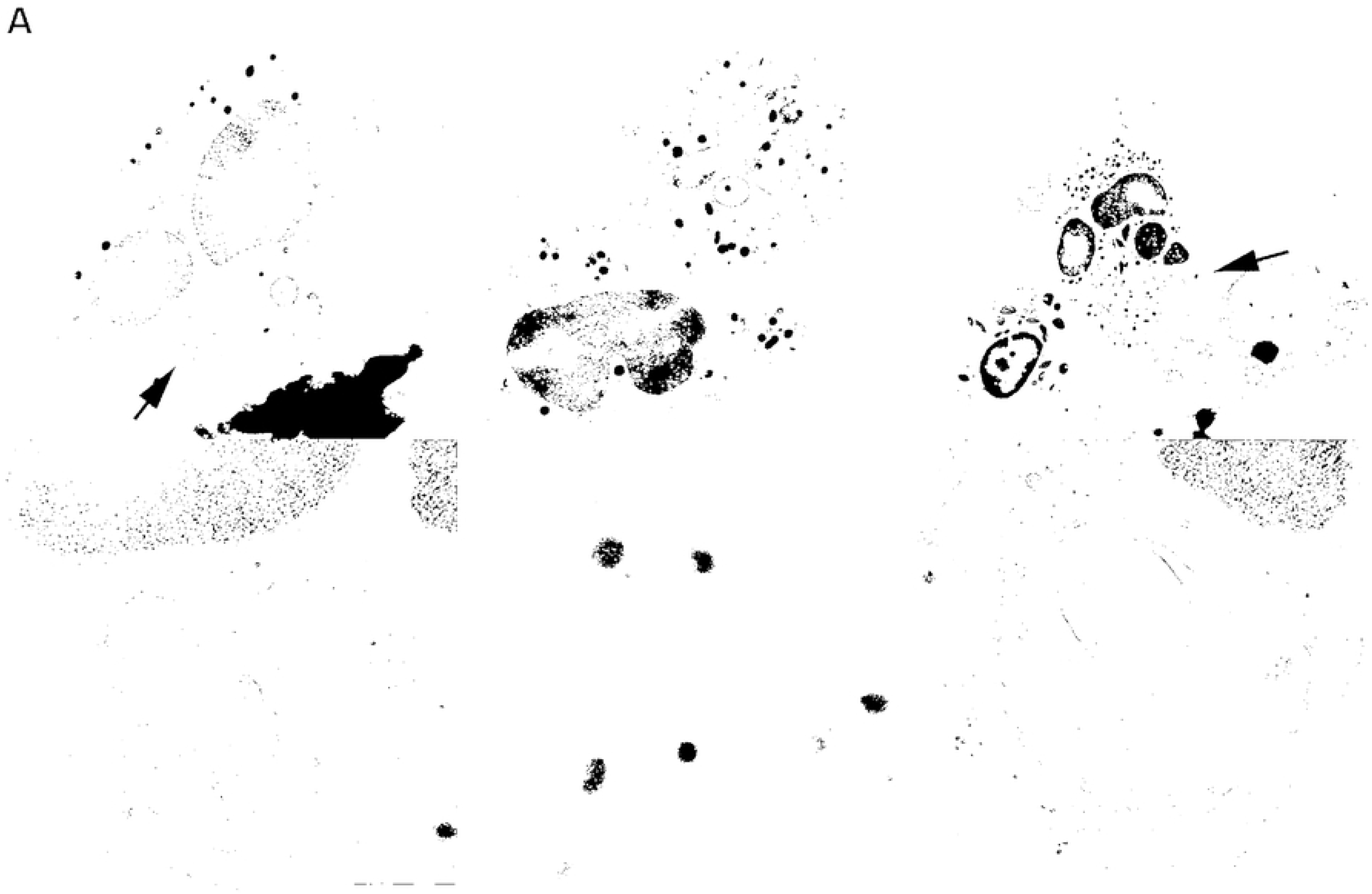
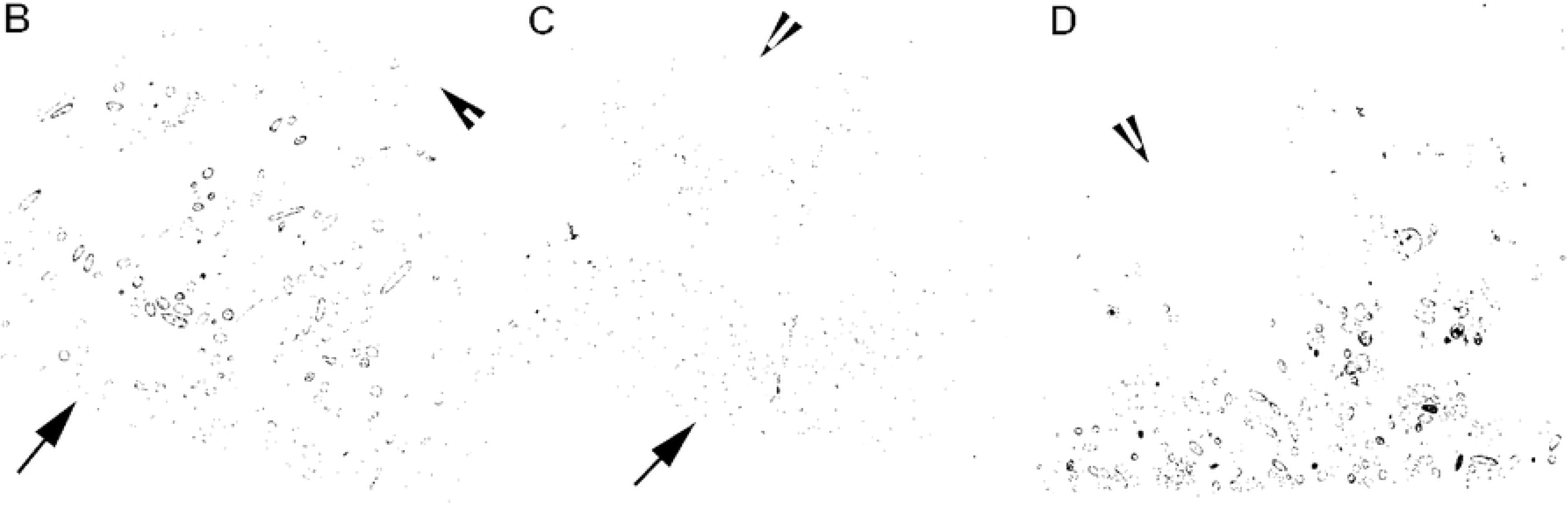
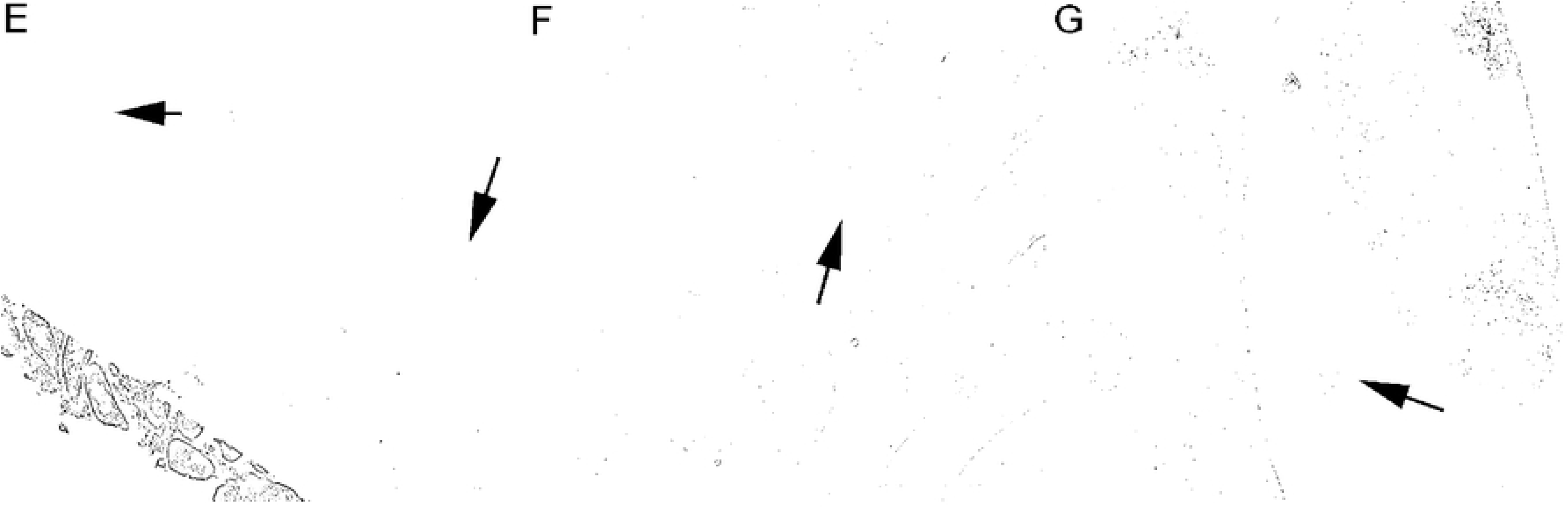
**A. *P. aeruginosa* is internalized by PMNs in the murine implant model.**Top row: TEM images showing intact and active PMNs containing internalized *P. aeruginosa* (arrows) 6h post-insertion of a pre-coated silicone implant (Bar: 2 μm). Bottom row: internalized bacteria are magnified (Bar: 500 nm). **B-D Interaction between PMNs and *P. aeruginosa* in the murine implant model.** Pre-coated silicone implants were inserted into the peritoneal cavity of BALB/c mice. The interaction between PMNs and bacteria was imaged by TEM at 6 h (B), 24 h (C), and 48 h (D) post-insertion. Black arrows: bacteria and biofilms, Arrow heads: PMNs (Bar: 5 μm). **E-G Biofilm formed by *P. aeruginosa* on a silicone implant at 24 h post-insertion.** TEM images showing *in vivo* biofilms at different magnifications. Matrix material (black arrows) can be seen between the bacteria. E) Bar: 2 μm; F) Bar: 1 μm; and G) Bar: 200 nm.

### *In vitro* interaction of *P. aeruginosa* and PMNs results in lysis of PMNs

When we examine the *in vitro* interaction between PMNs and aggregated *P. aeruginosa* microscopically, time series reveal that PMNs lyse resulting in release, expansion and dissolution of DNA (Movie 1). The staining of the nuclear content, DNA, with propidium iodide (PI), increases over time as the membrane of the PMNs becomes damaged. At the end the DNA disappears (the red color from PI fades) (Movie 1).

### PMN-derived eDNA surrounds *in vivo* biofilms in a murine implant model

To examine eDNA *in vivo*, we stained silicone implants *ex vivo* from the murine implant model with SYTO9 (green) which binds to DNA together with the Click-iT® DNA labeling technology kit to determine the origin (i.e., murine or bacterial) of the eDNA in biofilms *in vivo*. This method utilizes a modified thymidine analogue, 5-ethynyl-2’-deoxyuridine (EdU), which is incorporated into DNA during active DNA synthesis in the S-phase of the eukaryotic cell cycle. *Ex vivo*, PMN DNA was detected by labeling with a fluorescent Alexa Fluor® dye (pink). Treatment of healthy mice showed the presence of EdU labeled PMNs in the blood 2-5 days after treatment. Since the release of PMNs into the blood increases during an infection [23], we decided to treat the mice two days pre-infection to make sure EdU labeled PMNs would reach the implant and biofilm post-infection. In addition to bacteria aggregated within biofilms (Fig 2 and 3), long strings of eDNA (green) were clearly visible in the PMN accumulations surrounding the biofilms, but not as part of the biofilm itself (Fig 2 and 3).

**Fig. 2.**
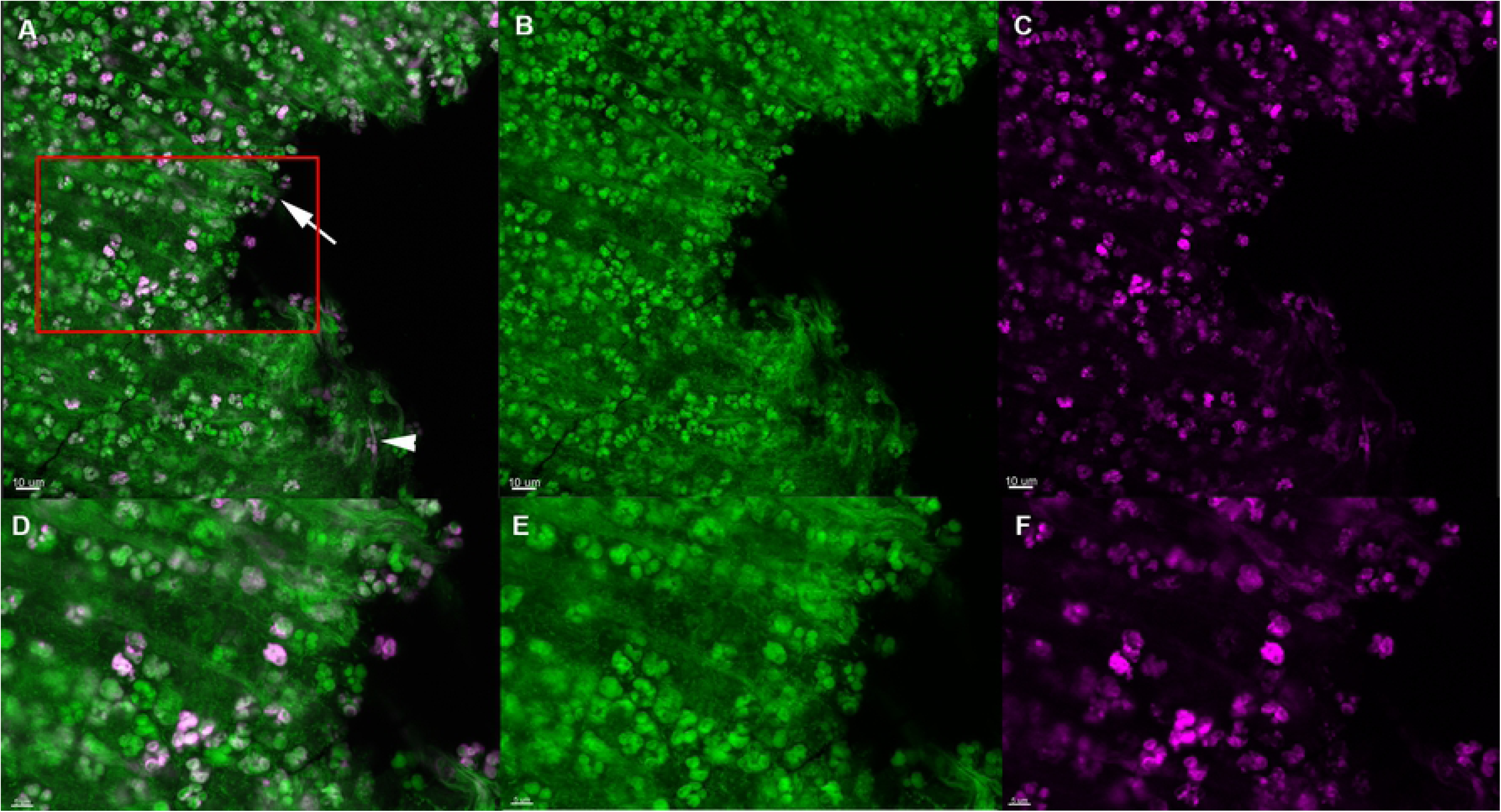
*In vivo* DNA labeling of PMNs in the murine implant model. PMNs in the murine implant model were labeled with Click-iT®, in which a modified thymidine analogue, EdU (5-ethynyl-2’-deoxyuridine), is incorporated into DNA during active DNA synthesis in murine immune cells *in vivo*, and hence will label only DNA originating from murine PMNs. DNA is labeled *ex vivo* with Alexa Fluor® 647 (pink) and counterstained with SYTO9 (green) on an implant 24h post-insertion in the peritoneal cavity. External to the biofilms SYTO9 (green)-stained DNA strings (white arrows) were also observed as well as a few pink DNA strings (white arrowheads). A, D) merged images showing both EDU (pink) and SYTO9 (green) staining. B, E) images showing only the SYTO9 staining. C, F) images showing only the EDU staining. Red square in A indicates magnified area in D-F.

**Fig. 3.**
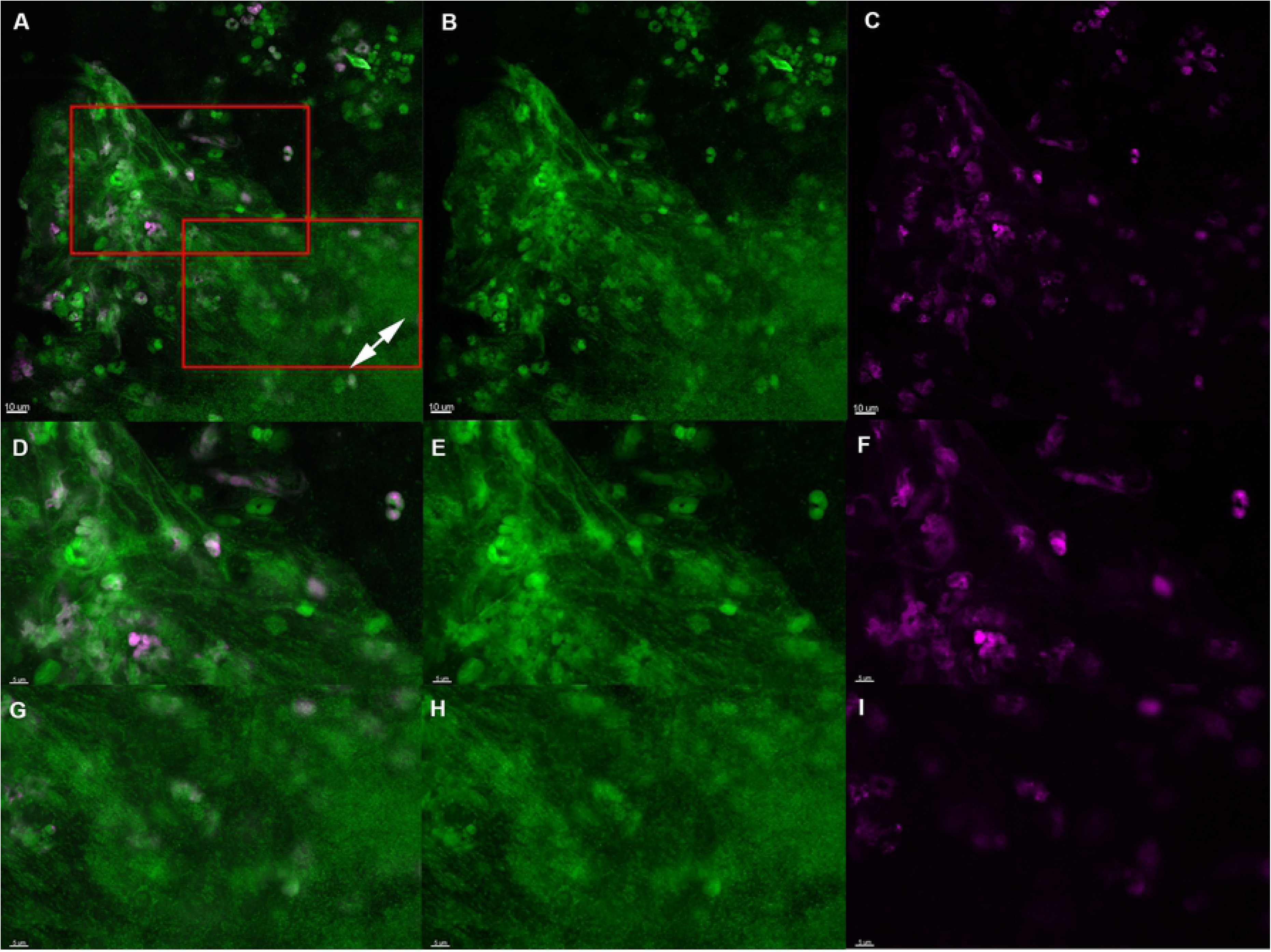
The *in vivo* biofilm lack PMN-derived DNA 24h post-insertion in the murine implant model. PMNs in the murine implant model were labeled with Click-iT®, in which a modified thymidine analogue, EdU (5-ethynyl-2’-deoxyuridine), is incorporated into DNA during active DNA synthesis in murine immune cells *in vivo*, and hence will label only DNA originating from murine PMNs. DNA is labeled *ex vivo* with Alexa Fluor® 647 (pink) and counterstained with SYTO9 (green) on an implant 24h post-insertion in the peritoneal cavity. Labeling was observed in PMNs (pink) but was absent from biofilms (double headed arrows), suggesting that PMNs are not a source of eDNA in biofilms. A, D) merged images showing both EDU (pink) and SYTO9 (green) staining. B, E) images showing only the SYTO9 staining. C, F) images showing only the EDU staining. Red squares in A indicates magnified area in D-F and G-I.

As seen in Fig 2, eDNA strings labeled with pink dye are clearly visible in close proximity to PMNs 24h post-insertion of the implant. However, no pink eDNA was detected in areas occupied by bacterial biofilms (Fig 3 and 4) at 24h or 48h post-insertion, indicating that PMN-derived eDNA is not present internally within these structures.

**Fig. 4.**
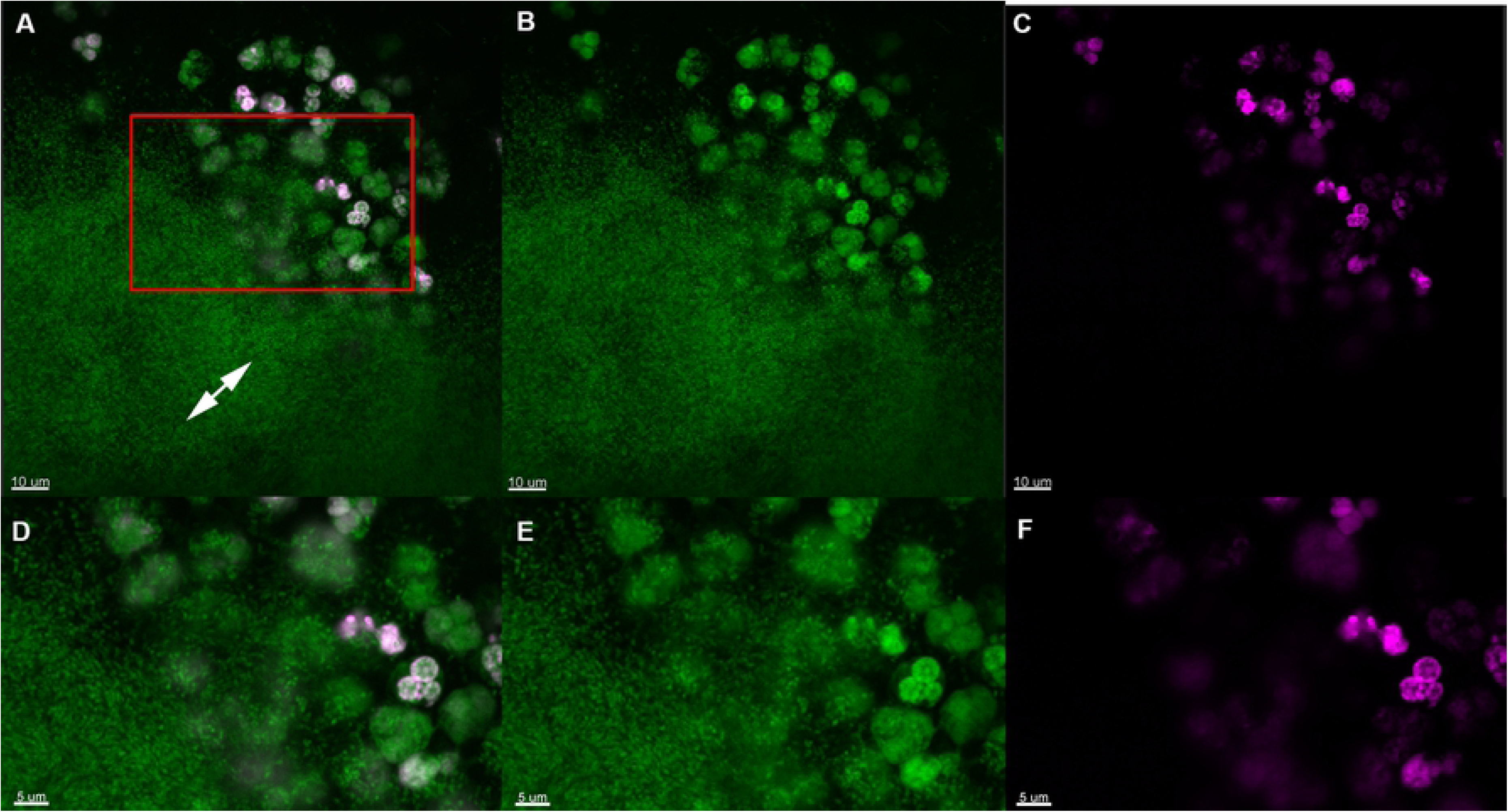
The *in vivo* biofilm lack PMN-derived DNA 48h post-insertion in the murine implant model. PMNs in the murine implant model were labeled with Click-iT®, in which a modified thymidine analogue, EdU (5-ethynyl-2’-deoxyuridine), is incorporated into DNA during active DNA synthesis in murine immune cells *in vivo*, and hence will label only DNA originating from murine PMNs. DNA is labeled *ex vivo* with Alexa Fluor® 647 (pink) and counterstained with SYTO9 (green) on an implant 48h post-insertion in the peritoneal cavity. Labeling was observed in PMNs (pink) but was absent from biofilms (double headed arrows), suggesting that PMNs are not a source of eDNA in biofilms. A, D) merged images showing both EDU (pink) and SYTO9 (green) staining. B, E) images showing only the SYTO9 staining. C, F) images showing only the EDU staining. Red square in A indicates magnified area in D-F.

### eDNA surrounds *in vivo* biofilms in lung tissue of chronically infected CF patients

Previously, peptide nuclear acid (PNA) fluorescence *in situ* hybridization (FISH) and 4’,6-diamidino-2-phenylindole (DAPI) have been used to identify bacterial aggregates surrounded by PMNs in paraffin-embedded sections of CF lung tissue [9]. The PNA probes bind to bacterial ribosomal 16S RNA [24] and DAPI binds to the A-T regions of double-stranded DNA. Here we used PNA FISH in combination with DAPI to show that eDNA (blue) was localized external to the biofilm in CF lung tissue (Fig 5). The biofilm stained red by a Texas Red-labeled *P. aeruginosa*-specific PNA probe (Fig 5A). The nuclei of PMNs were clearly stained, as were the strings of DNA outside the biofilm (Fig 5B).

**Fig. 5.**
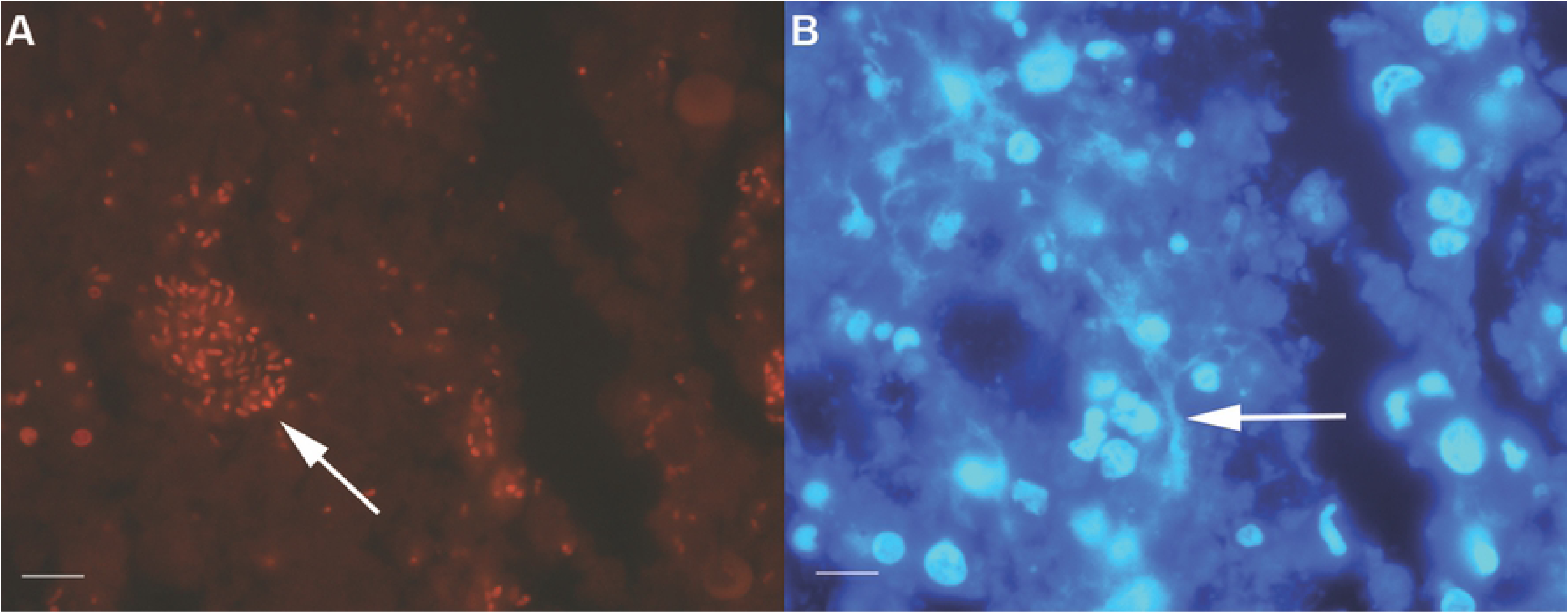
PNA FISH and DAPI staining of CF lung tissue. Aggregated *P. aeruginosa* (stained with PNA FISH (red)) (white arrows)) and eDNA (stained blue with DAPI) in CF lung tissue (A, B) Strings of DNA (B) are indicated by white arrows (Bar: 9 μm).

### Localization of histone H3 (H3), citrullinated H3 (citH3), and NE in a murine implant model

To localize components originating from murine PMNs during a bacterial biofilm infection, we stained paraffin-embedded sections of silicone implants isolated 24 h post-insertion in the peritoneal cavity of mice with antibodies specific for citH3 (Fig 6), H3 (Fig 7), or NE (Fig 8). All of these components, especially citH3, have also been associated with NETs. Hence, we used citH3 as a marker for NETs [25-28].

**Fig. 6.**
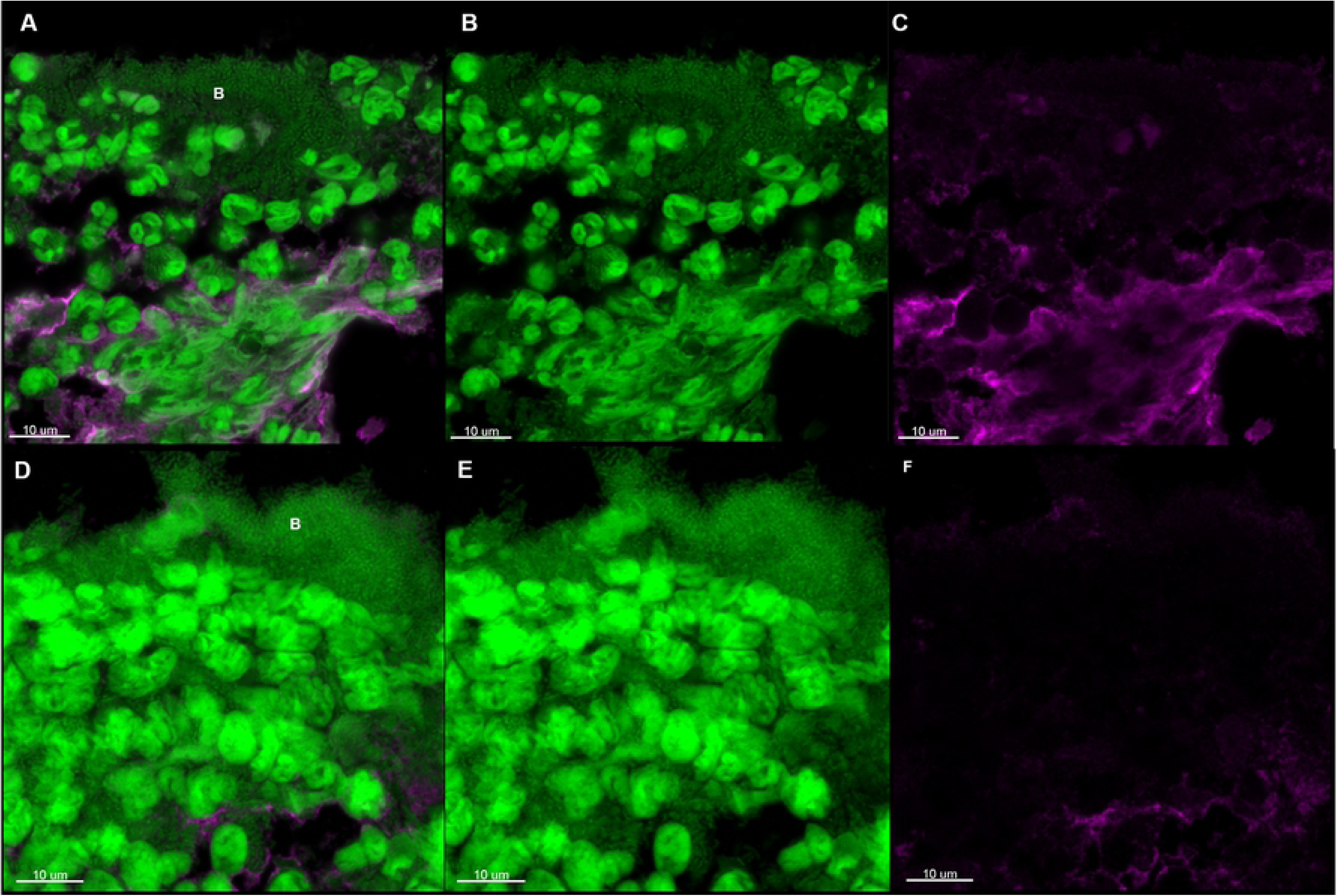
citH3 antibody staining of paraffin-embedded sections from the murine implant model. CSLM images showing paraffin-embedded sections of silicone implants from the murine implant model 24h post-insertion. The sections were stained with primary antibodies specific for citrullinated histone H3 (citH3). The secondary antibody was conjugated to Alexa Fluor 647 (pink). SYTO9 (green) was used as a counterstain. SYTO9 stains DNA in bacteria and eukaryotic cells. A, D) merged images of both the antibody (pink) and SYTO9 (green) staining. B, E) is only the SYTO9 staining. C, F) is only the antibody staining. The images represent two implants. The letter B indicates biofilm.

**Fig. 7.**
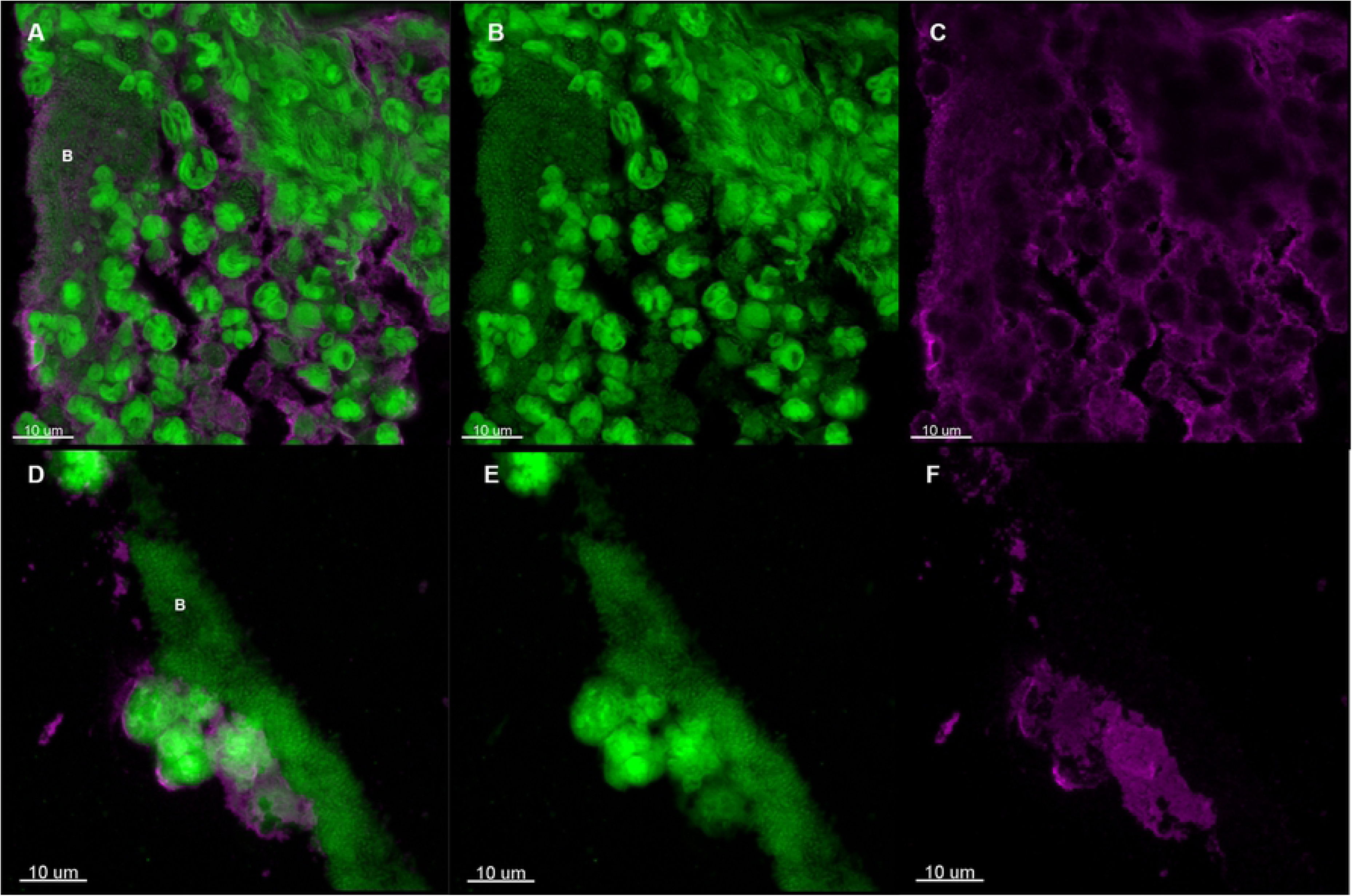
H3 antibody staining of paraffin-embedded sections from the murine implant model. CSLM images showing paraffin-embedded sections of silicone implants from the murine implant model 24h post-insertion. The sections were stained with primary antibodies specific for histone H3 (H3). The secondary antibody was conjugated to Alexa Fluor 647 (pink). SYTO9 (green) was used as a counterstain. SYTO9 stains DNA in bacteria and eukaryotic cells. A, D) merged images of both the antibody (pink) and SYTO9 (green) staining. B, E) is only the SYTO9 staining. C, F) is only the antibody staining. The images represent two implants. The letter B indicates biofilm.

**Fig. 8.**
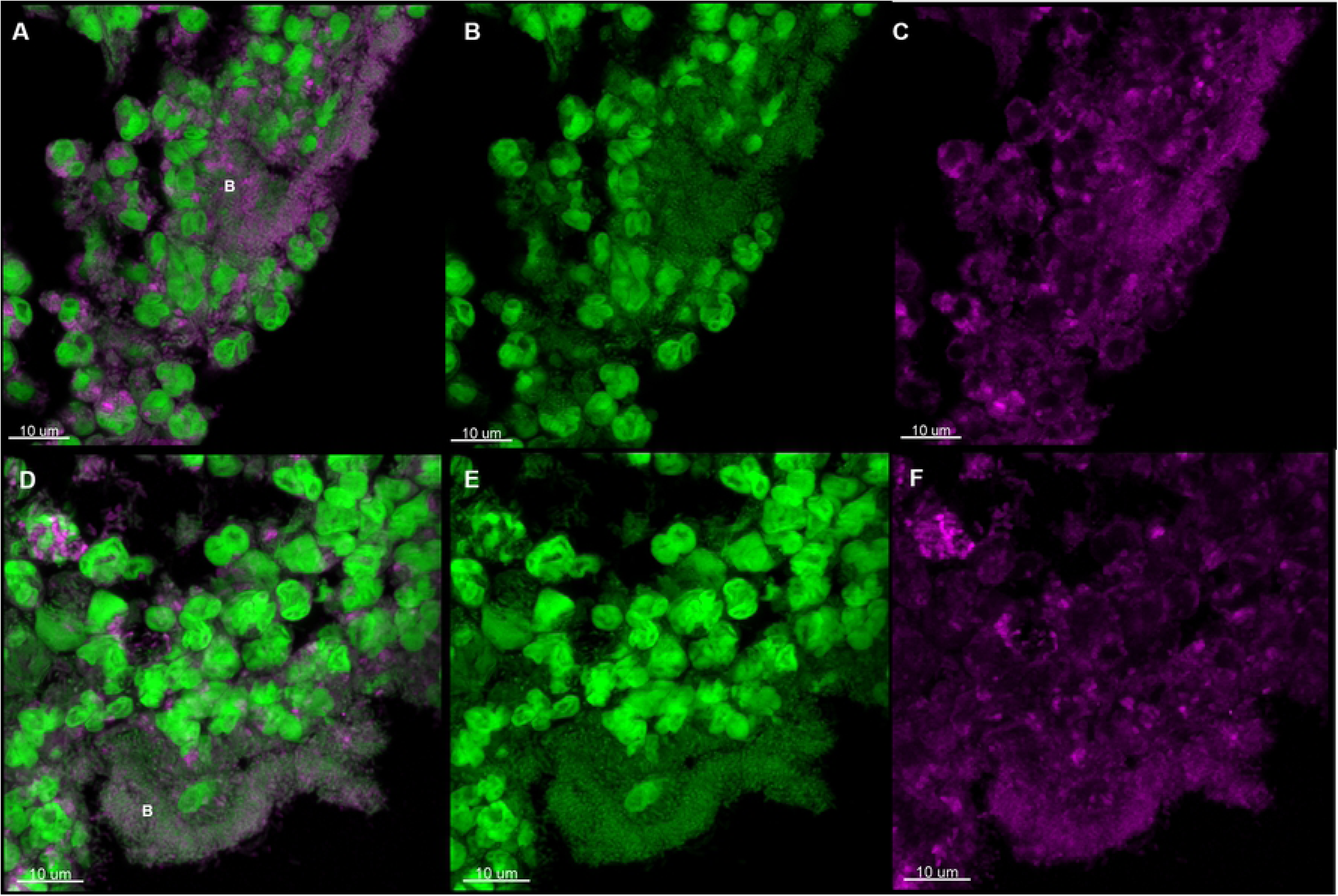
NE antibody staining of paraffin-embedded sections from the murine implant model. CSLM images showing paraffin-embedded sections of silicone implants from the murine implant model 24h post-insertion. The sections were stained with primary antibodies specific for neutrophil elastase (NE). The secondary antibody was conjugated to Alexa Fluor 647 (pink). SYTO9 (green) was used as a counterstain. SYTO9 stains DNA in bacteria and eukaryotic cells. A, D) merged images of both the antibody (pink) and SYTO9 (green) staining. B, E) is only the SYTO9 staining. C, F) is only the antibody staining. The images represent two implants. The letter B indicates biofilm.

H3 localized to areas surrounding bacterial biofilms. Only weak staining was observed inside the biofilm; these findings are in agreement with those described above (i.e., no eDNA originating from PMNs was observed within biofilms). citH3 localized with some PMNs, but the lack of citH3 signifies that the eDNA is not generated by NETosis. In contrast staining with specific antibodies against NE revealed distinct co-localization of biofilms and NE, indicating that NE is incorporated into biofilms (Fig 8). Control images are presented in Fig S4A.

### Localization of H3, citH3, and NE in lung tissue of chronically infected CF patients

Co-localization of NE and H3 with *P. aeruginosa* biofilms in the murine implant model led us to examine paraffin-embedded lung sections from explanted CF patients with chronic bacterial infections. To verify function and correct binding of the three antibodies when using human PMNs, we stained *in vitro* phorbol myristate acetate (PMA) generated NETs from human PMNs. We were able to show co-localization of NE and citH3 with NETs, but not with H3 (Fig S2). The lack of H3 co-localization correlates well with the fact that NETosis results in a partial degradation of H3 and posttranslational histone modifications and when using PMA as stimulation a total loss of full length H3 is observed [20, 29]. However, we did find co-localization of H3 with the nucleus of unstimulated PMNs, which indicates a functional antibody (Fig S3). NE also co-localized with the cytoplasma of unstimulated PMNs of paraffin-embedded sections (Fig S3).

In CF lungs the distribution of citH3, H3 and NE was similar to that observed in the murine implant model. CitH3 was observed in a few PMNs but did not co-localize with bacterial aggregates (Fig 9A-C). H3 was observed at the periphery of biofilms but not co-localized with the biofilm (Fig 9D-F). NE co-localized with bacteria within the biofilm (Fig 9G-I). Control images are shown in Fig. S4B. To estimate the co-localization of NE, H3 and citH3 with *in vivo* biofilms we used Manders’ co-localization coefficient as seen in Fig 9J. A value of 1, means that the green pixels (SYTO9) are perfectly co-localized with the pink pixels (antibody). The co-localization coefficient confirmed what we visually observed, H3 localized outside of biofilms (p<0.0001), citH3 sometimes to outside of biofilms (p<0.0007) and NE localized primarily to biofilms (p<0.25). There was not a significant difference between the outside and biofilm areas for NE. Some of the biofilms seemed to be located in a substance material, presumably alginate as previously shown [9] due to mucoid *P. aeruginosa*. These biofilms did not co-localize with NE. Instead NE was co-localized with the surrounding PMNs.

**Fig. 9.**
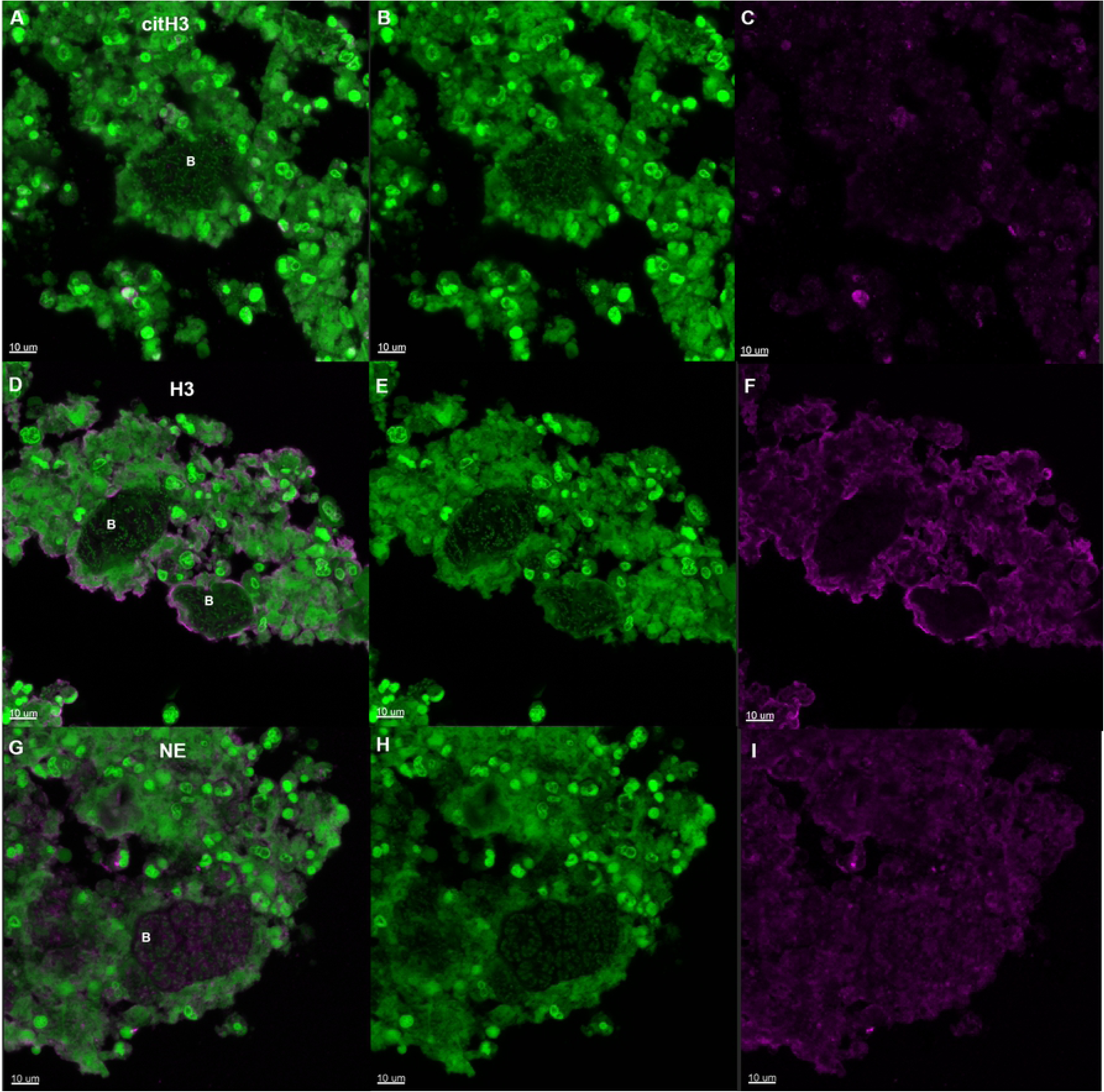
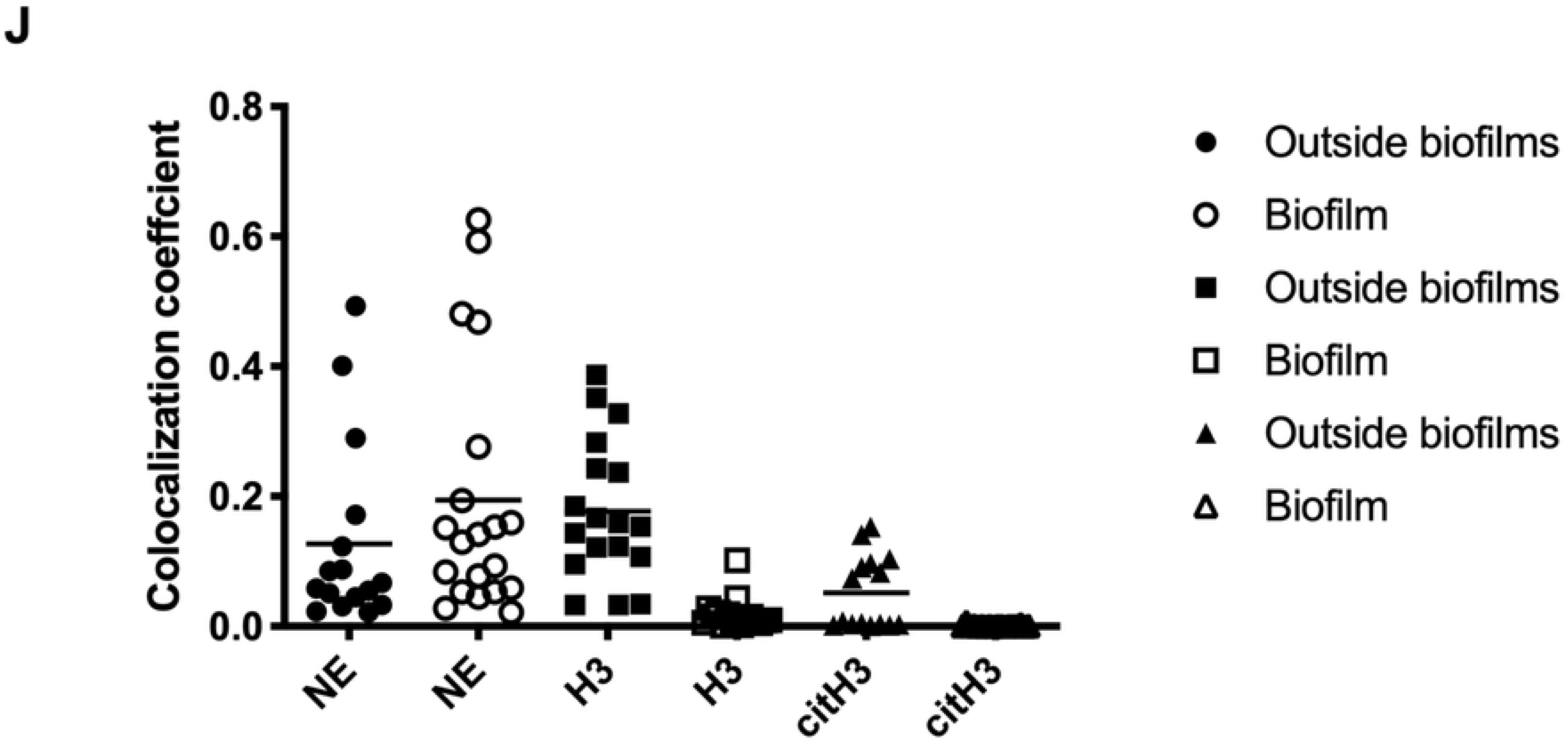
Immunostaining of paraffin-embedded sections from the CF lung tissue. CSLM images showing paraffin-embedded sections of CF lung tissue. The sections were stained with primary antibodies specific for citrullinated histone H3 (citH3) A-C). Histone H3 (D-F) or neutrophil elastase (NE) (G-I). The secondary antibody was conjugated to Alexa Fluor 647 (pink). SYTO9 (green) was used as a counterstain. SYTO9 stains DNA in bacteria and eukaryotic cells. A, D, G) merged images of both the antibody (pink) and SYTO9 (green) staining. B, E, H) is only the SYTO9 staining. C, F, I) is only the antibody staining. The images represent samples from three different CF lungs. B indicates biofilm. J: Co-localization of biofilm with antibodies in CF lungs shown as Manders’ co-localization coefficient. “Outside biofilm” are areas containing PMNs and “biofilms” are ROI defined biofilms. Sample size (n): NE outside biofilms (16), NE biofilms (20), H3 outside biofilms (18), H3 biofilms (17), citH3 outside biofilms (15), citH3 biofilms (18). There was significant difference between H3 outside biofilms and H3 biofilms (p<0.0001) and citH3 outside biofilms and biofilms (p<0.0007). No significant difference was found between NE outside biofilms and biofilms (p<0.25). p<0.05 was considered significant.

## Discussion

Surprisingly, our findings show that, *in vivo*, eDNA is concentrated external to bacterial aggregates rather than inside as shown previously *in vitro* by us and others [1, 4]. In fact, the lack of SYTO9 and DAPI staining within the bacterial aggregates in chronic biofilm infections truly questions the significance of the, otherwise *in vitro* important, role of eDNA in biofilms *in vivo* as a scaffold of the biofilm. Instead it highlights the important and detrimental feature of host-derived eDNA as a protective shell surrounding the bacteria. This barrier of host eDNA surrounding a biofilm *in vivo*, which could both limit dissemination of bacteria and shield the biofilm from phagocytosis, may be manifested by necrotic lysis of PMNs rather than through NETosis. Thus, our results are in support of deposition of PMN content in a zone between the PMNs and the biofilm as recently demonstrated in experimental *P. aeruginosa* ocular biofilm infection where the deposition of NETosis derived material prevented dissemination [30]. In our study, we found only scarce amount of host-derived eDNA and citH3 suggesting a modest contribution by NETosis to the material in the zone between PMNs and biofilm. However, dissemination of the bacteria may be prevented by the NE localized to the biofilm and therefore we propose other mechanisms than NETosis to contribute to the stalemate and chronicity of biofilm infection as illustrated in Fig. 10.

**Fig. 10.**
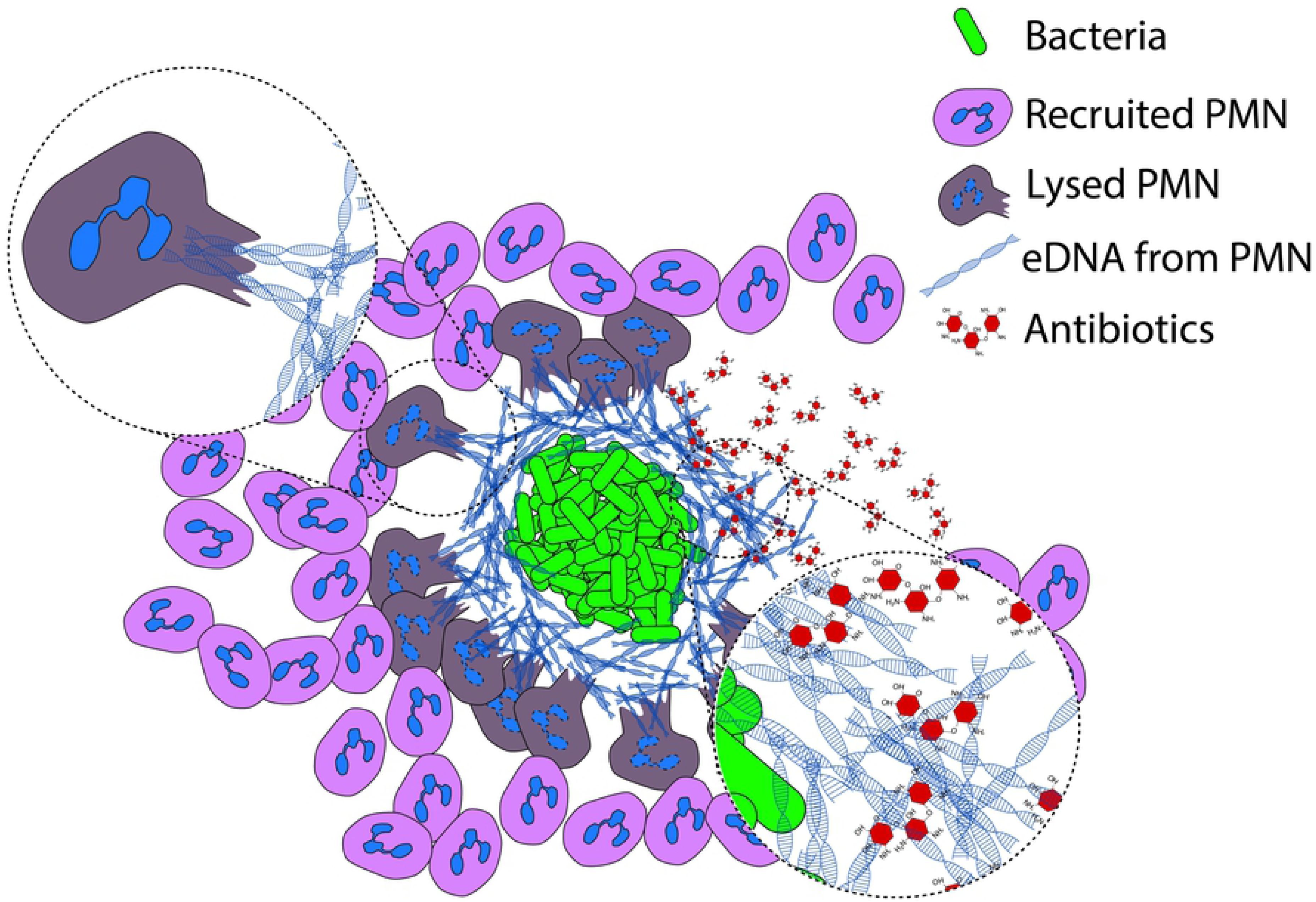
Hypothesis for formation of an eDNA shield in chronic bacterial infections *in vivo*. Early during biofilm formation, PMNs are able to phagocytose and destroy single bacteria or very small particles of bacterial cells. As the biofilm develops, bacterial aggregates evade phagocytosis and induce a necrotic cell death of PMNs. In chronic bacterial infections PMNs are continuously recruited to the site of biofilms where they release eDNA via necrosis. This released eDNA does not become incorporated into the biofilm itself. The PMN-derived layer of eDNA, which constitutes a secondary matrix, may provide a passive physical shield for the biofilm against cationic antibiotics such as tobramycin and additional phagocytes.

Using TEM we observed biofilm formation and swollen PMNs with damaged membranes, which is in agreement with our former SEM observations [22] showing PMNs failing to eradicate bacterial aggregates. The matrix material seen between the bacteria in the biofilm has not been fully characterized, but *in vitro* studies show the importance of eDNA for initial formation of young surface biofilms *in vitro* [4]; accordingly, it is believed to be an important structural component [6, 8]. Furthermore, *P. aeruginosa* has been shown to be attached to eDNA in CF sputum [31] and incorporate external eDNA into the biofilm matrix *in vitro* [5]. PMNs were enlarged and had damaged cell membranes, presumably due to exposure to the bacterial virulence factor, rhamnolipid, which is detrimental to PMNs [32, 33]. It has previously been shown that *P. aeruginosa* produce rhamnolipids which cause PMNs to swell and burst, thereby releasing their cytoplasmic and nuclear content into the surrounding area [34]. These cellular components including eDNA from necrotic PMNs are believed to serve as a biological matrix for biofilm formation and a protective shield within the biofilm increasing tolerance to treatment to tobramycin *in vitro* [5, 31]. This was confirmed in a mouse model where a *rhlA* mutant (unable to excrete rhamnolipid) did not cause any PMN lysis and no release of cytoplasmic and nuclear content, thus consistent with a mechanism of lysis by the biosurfactant rather than a programmed NET pathway for eDNA release [32].

The lack of strings in *in vivo* biofilms indicate that unlike *in vitro* biofilms, eDNA is not a direct part of the biofilm. Visually eDNA from *P. aeruginosa* has been shown to form a net both in a stationary grown biofilm and in the flow cell biofilm [1, 8]. Furthermore, eDNA from *P. aeruginosa* is believed to be double-stranded DNA [8] and since DAPI bind to double-stranded DNA the lack of DAPI staining signifies the lack of eDNA in the *in vivo* biofilm. Also, the measurement of double-stranded DNA has been shown to decrease when *P. aeruginosa* change from the planktonic state to the aggregating state [35]. However, the eDNA of a plate grown colony of *P. aeruginosa* have been shown to be 5 μm long [36] which mean that visually the eDNA fragment would be almost as small as a single *P. aeruginosa* bacterium. Visually this does not correlate to our *in vivo* and others *in vitro* findings. However, we might have to rethink the need for a structural matrix depending on the environment of the biofilm outside the laboratory. Maybe when the biofilm is exposed to high forces such as fluid it needs to establish a solid structure, but in the CF lung another type of structure is established within the mucus to withstand the immune cells. Accordingly, this indicates that neither bacterial nor PMN-derived eDNA is incorporated into biofilms *in vivo*, but eDNA from necrotic PMNs is more likely accumulating outside the biofilms.

Since we only found host-derived eDNA outside infectious biofilms *in vivo*, we speculated whether *in vivo* biofilm matrix structures comprised other components originating from PMNs (e.g., H3, citH3, or NE). PMNs undergoing apoptosis, necrotic cell death, or NETosis release different histones (i.e., H3 during necrotic cell death and citH3 during NETosis), either into the surrounding area or attached to NETs [17]. Release of these extracellular histones promotes secretion of pro-inflammatory cytokines and recruitment of immune cells to sites of infection [37]. Histones bind to lipopolysaccharide (LPS) derived from *P. aeruginosa*, which neutralizes the positive charge of the histone molecule [38]; with the outcome of antimicrobial activity [39]. The fact that histones are antimicrobial [17, 40], might not be the case in this study. Since H3 is rich in arginine it may rather play a role in the arginine pathway in *P. aeruginosa*, which breaks down arginine to ornithine via the arginine deiminase pathway to sustain growth under anoxic conditions. Ornithine is a carbon source used to maintain growth during periods in which nitrate is either limited or lacking [41-43]. Arginine has also been shown to contribute to biofilm formation [44].

Here, we showed that H3 is mostly present along the periphery of biofilm aggregates, but not within the biofilms. This could indicate that H3 is still bound to eDNA. By contrast, citH3 did not localize with biofilms in either the mouse model or CF lung samples. CitH3 is a post-translational modification of H3. Conversion of arginine to citrulline is accomplished by peptidylarginine deiminase 4 (PAD4) and is a prerequisite for *in vitro* generated NETosis since the modification of arginine residues to citrulline changes the charge of the histones, leading to massive chromatin de-condensation, NETs [26, 27, 45]. NETs are highly de-condensed chromatin structures that comprise mainly DNA, although histones and antimicrobial granular substances such as NE are also present [17]. Histone citrullination *in vitro* depends on stimulation of PMNs [46, 47]; indeed, an increase in histone citrullination is associated with chromatin de-condensation during NETs formation and LPS-induced early responses to inflammatory stimuli of PMNs [25, 26]. However, one widely used stimulus, PMA, has resulted in conflicting results in the literature and has been shown to induce NETs with and without co-localized citH3 and H3 [17, 47, 48]. In addition citrullination of H3 is strongly reduced when PMNs are stimulated during anoxic conditions [49], which is present in the endobronchial secretion in infected CF lungs [50, 51].

The lack of host-derived eDNA and citH3 associated with bacterial aggregates in the present study demonstrates that NETosis probably does not occur in chronic biofilm infections. Release of DNA under these conditions is not an active process as in NETosis; rather, it is more likely due to lysis of PMNs [34] which correlates to the presence of H3 and absence of citH3. In addition, our findings that NETosis seemingly plays no role in chronic *P. aeruginosa* infections correlates to the depletion of molecular oxygen at the site of chronic biofilm infections [52], thereby preventing NADPH oxidase from generating the reactive oxygen species (ROS) needed to release NETs [53]. Thus, PAD4-dependent NET formation might not be responsible for DNA release in CF airways and we speculate whether citH3 may serve as a biomarker for discriminating chronic from acute infections. However, NET generation independent of ROS generated by NADPH oxidase has been shown *in vivo* [54] therefore further work is needed to confirm this. Nevertheless, eDNA in sputum samples from CF patients have been shown to originate from PMNs [55] with NETs as the major component [56]. However, since NETs are antimicrobial this should benefit the CF patients since this should lead to eradication of the present bacteria. On the contrary eDNA has been shown to increase the viscosity of the mucus and bind antibiotics such as tobramycin [57, 58]. Furthermore, by treating the CF patients with DNase an improved lung function in CF patients and an increased effect of antibiotics have been reported [59, 60].

We did observe NE within biofilms, although this may have been derived from a necrotic cell death of PMNs external to the biofilm, followed by passive diffusion into the aggregate structure. NE is stored in primary (azurophilic) granules and released upon degranulation, phagocytosis, and cell death. NE cleaves a wide variety of proteins, including bacterial virulence factors [61]. The purpose of NE is to degrade phagocytosed proteins; however, in CF patients NE damages the airways [62]. Furthermore, NE is an important component of NETs [17]. However, since NE was found within the biofilms and eDNA was not supports the lack of NETs, since in these structures NE is bound to NETs. Furthermore, unlike in acute infections caused by

Gram-negative bacteria *P. aeruginosa* is not killed by NE following aggregation and biofilm formation [63]. The absence of NE-mediated killing is likely attributed to production of the protease inhibitor ecotin by bacteria [64]. In addition, NET formation induced by *P. aeruginosa* depends strongly on flagella motility [65], which may further explain the absence of NETs in CF lungs in which the flagella of *P. aeruginosa* are down-regulated [66] and in which flagellin, which is the major component of flagella, may be cleaved by NE [67]. Since a decrease and/or absence of flagella motility is associated with formation of bacterial biofilms [68], a lack of motile flagella may lead to a lack of detectable NETs.

In conclusion, this study is the first to examine the direct distribution of host and bacterial eDNA during chronic bacterial infections *in vivo*. Unlike after stimulation *in vitro*, stimulation of PMNs by bacterial aggregates during chronic infections *in vivo* does not seem to trigger active release of NETs; however, necrotic PMNs do release eDNA, histone H3, and antibacterial enzymes such as NE.

We did not find evidence of eDNA being part of *in vivo* biofilms, instead the PMN-derived layer of eDNA, which constitutes a secondary matrix, may provide a passive physical shield for the biofilm against cationic antibiotics such as tobramycin and additional phagocytes. A similar spatial arrangement has recently been described in a murine model of *P. aeruginosa* infection of the cornea in which the bacterial biofilm and neutrophil layers are separated by a “dead zone” rich in eDNA and NE [69].

Thus, we believe that eDNA released by PMNs due to a necrotic cell death contribute to the protection afforded to bacteria by biofilms (Fig. 10). These novel findings shed new light on the origin and role of eDNA in *in vivo* biofilms and unexpectedly question the role of extracellular host DNA during inflammation associated with chronic bacterial infections and of NET activation.

## Materials and Methods

### Murine implant model

All experiments were performed using a wild-type (WT) *P. aeruginosa* strain obtained from Professor Barbara Iglewski (University of Rochester Medical Center, NY, USA). The strain is QS proficient, except for the reduced production of C4-HSL previously noted for this *P. aeruginosa* variant [70]. Bacteria obtained from freezer stocks were plated onto blue agar plates (State Serum Institute, Denmark) and incubated overnight at 37°C. Blue agar plates are used to select Gram-negative bacilli [71]. One colony was used to inoculate overnight cultures grown in Luria-Bertani (LB) medium at 37°C with shaking at 180 rpm. Female BALB/c mice (6 weeks-of-age) were purchased from Taconic M&B A/S (Ry, Denmark) and maintained on standard mouse chow and water *ad libitum* for 2 weeks before challenge. The murine implant model was generated as described previously [22, 72], with some modifications. Briefly, the silicon tubes (Ole Dich Instrumentmakers Aps. Silicone Pumpeslange 60 Shore A (ID: 4.0 mm, OD: 6.0 mm, wall: 1.0 mm) were cut to a length of 4 mm, and the bacterial pellet from a centrifuged overnight culture was resuspended in 0.9% NaCl (OD_600nm_ = 0.1). Mice were anesthetized by subcutaneous (s.c.) injection of hypnorm/midazolam (Roche) [hypnorm (0.315 mg ml^-1^ fentanyl citrate; 10 mg ml^-1^ fluanisone); midazolam (5 mg ml^-1^); and sterile water; 1:1:2 v/v]. The pre-coated silicone implants were inserted in the peritoneal cavity of the mice. The mice were treated with bupivacaine and Temgesic® as post-operative pain relief. Experiments were terminated with euthanization of the mice with intraperitoneal injection of pentobarbital (DAK; 10.0 ml kg^-1^ body weight).

### Preparation of samples for TEM

The silicone implants were fixed in 2% glutaraldehyde in 0.05 M sodium phosphate buffer pH 7.4. Growing biofilms on a silicone tube presents a challenge with respect to sample preparation prior to the different microscopy techniques. First, a traditional preparation method was used for TEM to get an indication of the interaction between PMNs and the biofilm. However, some of the steps involved in the embedding process were shortened. Samples were observed with a CM 100 TEM (Philips, the Nederlands) operated at 80kV. Imaged were recorded with a side-mounted Olympus Veleta camera (resolution, 2048 × 2048 pixels (2K × 2K)) and the iTEM (Olympus, Germany) software package.

### *In vitro* experiments with PMNs and *P. aeruginosa*

Wild-type *P. aeruginosa* (strain PAO1) was obtained from the Pseudomonas Genetic Stock Center (www.pseudomonas.med.ecu.edu; strain PAO0001). Bacteria were tagged with a stable green fluorescent protein (GFP) constitutively expressed by plasmid pMRP9 [73]. PAO1 cultures were grown for 24 h at 37°C in 12 well microtiter plates containing LB medium. Human blood was collected from healthy volunteers with the approval of the Danish Scientific Ethical Board (H-3-2011-117). PMNs were isolated as described [74]. Isolated PMNs were resuspended in krebs ringer buffer containing 10mM glucose at 37°C (final density, 2.5 × 10^7^ PMNs/ml). Propidium iodide (2.5 ug/ml) was used as an indicator of cell death. PAO1 (350 μl) and PMNs (50 μl) were mixed in a 12 well microtiter plate and observed under a confocal microscope for 35 min.

### Click-iT™

Mice were treated with EdU (Click-iT™ EdU Alexa Fluor™ 647 Imaging Kit) (C10640, ThermoFisher) for 2 days pre-infection and until termination of the experiment. Each mouse was injected i.p. twice a day with 25 mg/g EdU dissolved in 0.9% NaCl. Control mice received 0.9% NaCl without EdU. *Ex vivo* implants were fluorescently labeled using the Click-iT kit, according to the manufacturer’s instructions, and stained with SYTO9 (S34934, ThermoFisher) prior to observation under a microscope.

### *In vitro* generated NETs

*In vitro* NETs were generated according to Vong et. al. [75] and Brinkmann et. al. [76] with modifications. PMNs were isolated according to Hu et. al. [77] using polymorphprep with modifications. Isolated PMNs were suspended in RPMI 1640 with glutamine and were seated in an ibidi slide coated with poly-L-lysine (80604, ibidi) for 30 min before stimulation with PMA for 3h. The PMNs were incubated at 37 degrees and 5% CO_2_. PMNs was washed with PBS and fixed in 4% PFA pH 7.2 for 15 min, washed in 0.025% Triton X-100 twice before proceeding with antibody staining as described for paraffin sections. Before imaging the channels were filled with Ibidi mounting medium (50001, ibidi).

### *Ex vivo* paraffin samples

Samples from the mouse model, the lungs of CF patients, and unactivated human PMNs were fixed in 4% formalin and paraffin-embedded according to standard protocols. Sections (4 μm thick) were cut using a standard microtome, fixed on glass slides, and stored at 4°C until required.

### Lung tissue from chronically infected CF patients

Explanted lungs tissue from 3 contemporary intensively treated *P. aeruginosa* infected CF patients were examined [9].

### PNA FISH

PNA FISH staining was performed as previously described [9, 78]. Texas Red-labeled *P. aeruginosa*-specific PNA probe both in hybridization solution (AdvanDx, Inc., Woburn, MA), was added to each section and hybridized in a PNA FISH workstation covered by a lid at 55°C for 90 min. The slides were washed for 30 min at 55°C in wash solution (AdvanDx) and counterstained with DAPI (D1306, ThermoFisher).

### Antibody staining of paraffin-embedded tissue sections

Sections were deparaffinized in 99.9% xylene (twice for 5 min each time) and subsequently hydrolyzed in 99.9% ethanol (twice for 5 min each time), followed by 96.5% ethanol (twice for 3 min each time). Finally, sections were rinsed in water (twice for 5 min each time). Heat-induced antigen retrieval was performed in citrate buffer, pH 6.0 (Sigma C9999) for 15 min in a domestic microwave oven. For unstimulated human PMNs a permabilization step was included using 0.1% Triton X-100 for 20 min. Samples were washed twice (5 min each time) in 0.025% triton X-100, drained, and blocked for 2 h at room temp in 10% goat serum in 1% BSA/PBS. Samples were drained and incubated for 16–18 h at 4°C with a primary antibody (Abcam Cat# ab1791, RRID:AB_302613, Abcam Cat# ab5103, RRID:AB_304752, or Abcam Cat# ab21595, RRID:AB_446409), diluted 1:200 in 2% goat serum/1% BSA in PBS). Negative controls were isotype IgG (Abcam Cat# ab27478, RRID:AB_2616600) or absence of primary antibody. Samples were drained and washed in 0.025% Triton X-100 (twice for 5 min each) with gentle agitation. Samples were then incubated (1 h at room temp in the dark) with a secondary antibody (Abcam Cat# ab150083, RRID:AB_2714032) diluted 1:1000 in 1% BSA/PBS. Finally, samples were washed 3 times in PBS (5 min each time) and counterstained with SYTO9 (S34934, ThermoFisher). ProLong^™^ gold antifade mountant (Thermofisher, P36934) was applied before adding the coverslip.

### Co-localization coefficient

Co-localization refers to the geometric co-distribution of two fluorescent labels or color channels. Co-localization was acquired in Zen (version 2.1 SP3 FP2 (black)) as described by Zeiss “Acquiring and analyzing data for co-localization experiments in AIM or Zen software”. The Manders’ co-localization coefficient [79] for SYTO9 (green channel) was used to show the co-localization of the antibody (pink channel) using 2D images. Perfect co-localization will give the value 1. The crosshair was set to 1003 for the green channel and 2500 for the pink channel based on single stained paraffin sections according to the manual by Zeiss. Sections from 3 different patients were used. Biofilm and PMN areas were selected by region of interest (ROIs) in each paraffin section. All biofilm areas in one image were included. GraphPad Prism was used for statistical analysis using unpaired t-test. P<0.05 was considered significant.

### Confocal scanning laser microscopy

All observations and image acquisition were performed using a confocal scanning laser microscope (LSM510, LSM 710, or LSM 880; Carl Zeiss GmbH, Germany). Images were obtained using 40× 63× or 100× oil objective lenses. Images were scanned at 405 nm (blue), 488 nm (green), and 633 nm (far red). Images of the biofilms were generated using Imaris software including the magnifications (version 8.4, Bitplane AG). Images were processed for display using Photoshop software (Adobe).

## Ethics Statement

All animal studies were carried out in accordance with the European convention and Directive for the Protection of Vertebrate Animals used for Experimental and Other Scientific Purposes, and with Danish law on animal experimentation. All experiments were authorized and approved by the National Animal Ethics Committee, Denmark (The Animal Experiments Inspectorate, dyreforsoegstilsynet.dk; permit numbers: 2012−15−2934−00677 and 2014−15−0201−00259).

Human blood was collected from healthy volunteers with the approval of the Danish Scientific Ethical Board (H-3-2011-117). All donors provided written informed consent.

Infected tissue was collected following approval by the Danish Scientific Ethical Board (KF465 01278432). The samples used were collected with written informed consent for a previous study [9], and only from adults. The samples were anonymized for this study.

## Acknowledgments

We acknowledge the Core Facility for Integrated Microscopy, Faculty of Health and Medical Sciences, University of Copenhagen, Zhila Nikrozi for preparing samples for TEM, Heidi Marie Paulsen for preparing paraffin sections, and Steen Seier Poulsen for helpful discussions regarding antibody staining procedures. M.A., M.A., and T.B. were funded by the Lundbeck Foundation. M.A. was also founded by Hørslev-Fonden and Torben and Alice Frimodt Fonden.

## Author contributions

M.A. conducted the mouse experiments and antibody staining experiments, and wrote and edited the manuscript.

M.A. undertook microscopic analysis of Click-It and edited the manuscript.

K.Q. assisted with TEM preparation and image recording, and edited the manuscript.

P.Ø.J. performed the *in vitro* PMN experiments, contributed to data interpretation and edited the manuscript.

K.N.K helped with the co-localization, helped with figure 10 and edited the manuscript

P.S. contributed to data interpretation and edited the manuscript

T.B. performed the *in vitro* PMN experiments and PNA FISH, contributed to data interpretation and edited the manuscript.

## Supporting Information Legends

### Movie 1

Interaction between isolated human polymorphonuclear leukocytes (PMNs) and aggregated *P. aeruginosa* for 35 min. *P. aeruginosa* was grown for 24h prior to exposure to PMNs. PMNs are bursting, probably due to production of rhamnolipid by *P. aeruginosa*. *P. aeruginosa* are tagged with GFP, and dead cells are stained with propidium iodide (PI).

**Fig. S1. TEM images 48h post-insertion in the murine implant model**

TEM images displaying *P. aeruginosa* biofilms and matrix material at 48 h post-insertion in the murine implant model. Matrix material is denoted by black arrows. Scale bars: A) 5 μm, B) 2 μm.

**Fig. S2 *in vitro* PMA generated NETs and antibody staining**

Isolated human PMNs were stimulated with PMA for 3h *in vitro* and stained with antibodies. A, B) antibody staining against NE (pink) (A is a merged image of both the antibody (pink) and SYTO9 (green) staining, B) is only the antibody staining), C) antibody staining against H3, no pink staining was observed. D, E) antibody staining against citH3 (D is a merged image of both the antibody and SYTO9 staining, E is only the antibody staining). F) antibody staining using isotype IgG as a control. SYTO9 was used as a counterstain (green). NE and citH3 (pink) were associated to DNA (SYTO9), but H3 was not. No pink was imaged in the control (isotype IgG).

**Fig. S3. Immunostaining of unactivated paraffin embedded human PMNs**

Unstimulated human PMNs in paraffin sections were stained with NE or H3 (pink). The DNA of the PMNs was stained with SYTO9 (green). NE co-localized with the cytoplasma of the PMNs (A, B) and H3 with the nucleus (C, D). A, C) are merged images of both the antibody (pink) and SYTO9 (green) staining, B, D) are only the antibody staining).

**Fig. S4. Controls for immunostaining**

Controls for immunostaining of paraffin-embedded sections from the murine implant model and cystic fibrosis (CF) lungs. The paraffin-embedded sections were stained either with either isotype control IgG or in the absence of primary antibody. The secondary antibody was conjugated to Alexa fluor 647 (pink). SYTO9 (green) was used as a counterstain. SYTO9 stains DNA in bacteria and eukaryotic cells. B indicates biofilm and arrow heads points to PMNs.

